# Using SMOG 2 to simulate complex biomolecular assemblies

**DOI:** 10.1101/371617

**Authors:** Mariana Levi, Prasad Bandarkar, Huan Yang, Ailun Wang, Udayan Mohanty, Jeffrey K. Noel, Paul C. Whitford

## Abstract

Over the last 20 years, the application of structure-based (Gō-like) models has ranged from protein folding with coarse-grained models to all-atom representations of large-scale molecular assemblies. While there are many variants that may be employed, the common feature of these models is that some (or all) of the stabilizing energetic interactions are defined based on knowledge of a particular experimentally-obtained conformation. With the generality of this approach, there was a need for a versatile computational platform for designing and implementing this class of models. To this end, the SMOG 2 software package provides an easy-to-use interface, where the user has full control of the model parameters. This software allows the user to edit XML-formatted files in order to provide definitions of new structure-based models. SMOG 2 reads these “template” files and maps the interactions onto specific structures, which are provided in PDB format. The force field files produced by SMOG 2 may then be used to perform simulations with a variety of popular molecular dynamics suites. In this chapter, we describe some of the key features of the SMOG 2 package, while providing examples and strategies for applying these techniques to complex (often large-scale) molecular assemblies, such as the ribosome.

## 1 Introduction

When performing a molecular dynamics (MD) simulation, the equations of motions are repeatedly integrated for a given Hamiltonian. In the context of molecular biophysics, there is a wide variety of Hamiltonians available, where each is typically described in terms of its potential energy function (i.e. the “model”, or “force field”). Available models differ in terms of both spatial resolution (e.g. all-atom, or coarse-grained) and energetic detail. In the present chapter, we will discuss one class of models, called “structure-based models”. The hallmark feature of structure-based models is that the user predefines the dominant potential energy minima, prior to performing a simulation. Typically, the minima are defined such that they correspond to experimentally-obtained (i.e. “native”) structures. Originally, this type of model was inspired by the principle of minimal frustration [1], which states that the interactions that dominate the energy landscape of protein folding are those formed in the native configuration. This led to the development of early coarse-grained (i.e. “Gō-like”) models [2, 3] of proteins, which were subsequently extended to all-atom resolution [4, 5].

While purely structure-based models have been widely used in the study of protein folding and function [6, 7], these models can also serve as a basis for exploring the role of non-specific energetic interactions. For example, protein-DNA interactions have been extensively studied through the use of structure-based models that include electrostatic effects [8, 9], and RNA-RNA electrostatics have been characterized in the context of the ribosome [10]. Beyond the inclusion of electrostatic interactions, one can envision a more general spectrum of models that range from purely structure-based (i.e. only native interactions are stabilizing) to completely non-specific (e.g. CHARMM[11], AMBER[12]) representations.

To provide a flexible platform for the construction of structure-based model variants, we recently released the SMOG 2 [13] software package. The principle that guided development of this software is that any user should be able to define/extend structure-based models without modifying source code. To enable this, SMOG 2 reads user-provided “template” files that define rules for constructing structure-based models. SMOG 2 maps the definitions in the template files onto a biomolecular system, which allows the user to perform their simulations with a number of popular MD engines (Gromacs [14], NAMD [15], or openMM [16]). This allows the user to rapidly alter the resolution of the model, introduce charges on specific atoms/residues, or simulate new residues (e.g. modified residues). From the user’s perspective, only simple modifications to XML-formatted template files are required to construct a structure-based model that is suited to address a specific physical questions.

In this chapter, we provide step-by-step descriptions of how to utilize the SMOG 2 software package. In particular, we provide examples for how to effectively utilize this software to simulate large biomolecular systems, such as the ribosome. Our intention is that this guide will provide the technical steps necessary for a typical computational researcher to develop their own structure-based models and apply them to complex molecular assemblies. For a thorough discussion of the general utility of these models, other references are recommended [17, 18].

## 2 Materials

This chapter describes the SMOG 2 software package [13]. The commands and modules are specific to a recently released version of SMOG 2 (v2.1). As mentioned above, the purpose of SMOG 2 is to read a set of template files, which define the rules for constructing a structure-based model, and then map these interactions onto a specific structure of a biomolecular system (provided in PDB format). The output of SMOG 2 is a set of force field and coordinate files that are formatted for use with Gromacs [14] (v4, or v5). Here, we provide instructions for how to utilize these models with Gromacs v5. However, other groups have ported these models to NAMD and openMM, thereby allowing the SMOG 2 output files to be used with a broader range of MD engines (see user’s manual for details).

## 3 Methods

In this section, we first provide an overview for how to generate a structure-based model and perform a simulation. This is followed by tutorials for using SMOG tools, as well as examples for how to design new models (e.g. novel residues, or introduction of charges.)

### 3.1 Basic steps when using SMOG 2

To illustrate the basic utility of SMOG 2, we will briefly describe the steps necessary to simulate an arbitrary protein. For this example, we will simulate chymotrypsin inhibitor 2 [19] (CI2). For the steps below, $SMOG_PATH will refer to the main directory of the locally-installed version of SMOG 2, and smog2 will refer to the SMOG 2 executable (located in $SMOG_PATH/bin). It is assumed that the SMOG executables are in the user’s path (this is automatically performed by the configure.smog2 script).

#### 3.1.1 Preprocess the structure (PDB) file

Download and save the the atomic coordinates of CI2 (PDB ID: 2CI2) to a file named 2CI2.pdb. The PDB identifiers ATOM, HETATM, BOND, TER, and END are recognized by SMOG 2. TER lines separate covalently bonded chains and BOND lines indicate that SMOG 2 should include system-specific covalent bonds, e.g. a disulfide bond. Since CI2 is a monomer with standard residue naming, only the ATOM and END lines are relevant. One may extract these lines with a simple one-line grep command:

~~~
>$ grep ’^ATOM\ |^END’ 2 CI2. pdb > CI2. atoms. pdb
~~~

Once you have removed all comments from the PDB file, it is necessary to rename any terminal residues, such that they are consistent with default SMOG 2 conventions. For example, a C-terminal protein residue has an OXT atom. In order for the template files to be properly mapped to the PDB structure, terminal residues must be distinguished from non-terminal residues. To facilitate this step, SMOG 2 includes the smog_adjustPDB tool. When using the default SMOG models, smog_adjustPDB may be used with the following flags:

~~~
>$ smog_adjustPDB *−* i CI2.atoms.pdb *−* default
~~~

If an output file name is not specified, then the new pdb file will be named adjusted.pdb. For tips on PDB formatting, see Note 1.

#### 3.1.2 Use SMOG 2 to generate a forcefield

For this example, we will use the all-atom structure-based model [5], which is defined by the template files found in the directory $SMOG_PATH/share/templates/SBM_AA. To generate the force field files, issue the command:

~~~
>$ smog2 *−* i adjusted.pdb *–*AA
~~~

An equivalent invocation would be:

~~~
>$ smog2 *−* i adjusted.pdb *−*t $SMOG_PATH/ share / templates /SBM_AA
~~~

Note that, if you want to use a non-standard (e.g. user-modified) force field, then the second invocation should be used, where the location of the template files would be given after the -t flag.

After SMOG 2 successfully completes, four files will be generated (default names below):

- **smog.gro**: Atomic coordinates in Gromacs format.
- **smog.top**: Topology file that defines the potential energy function.
- **smog.contacts**: List of atom pairs used to define stabilizing “native” contacts in smog.top (listed in the [pairs] section).
- **smog.ndx**: Gromacs index file listing the atoms of each chain as a separate group.

In this example, the file 2CI2.pdb works smoothly with SMOG 2 since it contains coordinates for all non-Hydrogen atoms, uses standard nomenclature for the 20 amino acids and their constituent atoms, and it does not have non-standard amino acids. PDB files often contain atoms, residues, or small molecules that differ from, or are not defined in, the SMOG 2 default templates. Such instances will trigger errors during SMOG 2 processing.

#### 3.1.3 Perform the simulation in Gromacs

After using SMOG 2, one may perform a simulation with a number of MD engines. For the steps below, we provide the commands necessary to utilize Gromacs v5. When using Gromacs, one must create a portable xdr file (in the example below, run.tpr) that contains the coordinates, force field and simulation parameters. The .mdp file defines the simulation settings, such as the integrator/thermostat (e.g. Berendsen, Langevin Dynamics, Nosé Hoover), time step size, number of time steps and simulated temperature. For the commands below, it is assumed that the Gromacs 5 executable (gmx) is in your path. To prepare your simulation, you should first center your molecule within a box using the editconf module of Gromacs:

~~~
>$ gmx editconf *−* f smog.gro *−*c *−*d 2 *−*ocentered.gro
~~~

In this example, -d 2 tells Gromacs to define a box such that the solute is 2 nm from any edge (For tips on setting the box size, see Note 2). Next, use the grompp module of Gromacs to prepare your tpr file:

~~~
>$ gmx grompp *−* f min.mdp *−*p smog.top *−*maxwarn 1 *−*ccentered.gro *−*o min
~~~

In this example, it is assumed that the mdp file named min.mdp be being used to instruct Gromacs to perform steepest-descent energy minimization (see user manual, or example files distributed with SMOG 2, for recommended settings in the mdp file). Note: It is typically necessary to use the flag -maxwarn 1 when using SMOG models. The reason is that, to improve performance, periodic boundary conditions are almost always employed. However, since most SMOG models do not include solvent, the potential energy is translationally and rotationally invariant. Accordingly, it is necessary to remove center of mass rotation, even though periodic boundaries are used. If the molecule does not traverse any boundaries (which can be ensured by using a very large box), one may ignore the warning regarding the removal of center of mass rotational freedom in a periodic system.

After you have created the .tpr file with grompp, perform energy minimization with mdrun:

~~~
>$ gmx mdrun *−*v *−*deffnm min *−*noddcheck [*−* nt numberOfThreads]
~~~

By using the flag -deffnm min all output files will be named “min”, with the appropriate suffix appended. After minimization, repeat the grompp and mdrun steps using an different mdp (for this example, run.mdp) file in which integrator = steep is replaced with integrator = sd. This will indicate to Gromacs that stochastic dynamics should be used.

~~~
>$ gmx grompp *−* f run.mdp *−*p smog.top *−*maxwarn 1 *−*c min. gro *−*o run.tpr
>$ gmx mdrun *−*v *−*deffnm run *−*noddcheck [*−* nt numberOfThreads]
~~~

### 3.2 Simulating a portion of a larger assembly

It is common practice in the MD community to only simulate a portion of a larger complex. For example, recent studies on the ribosome have involved free-energy calculations where only the atoms near bimolecular interfaces are represented [20, 21]. Large-scale conformational changes of tRNA inside of the ribosome have also performed where only a portion of the ribosome is explicitly represented [10, 22]. In all of these examples, the rationale for simulating a subset of atoms is that the bimolecular interactions of interest are not always associated with global conformational rearrangements. Accordingly, to reduce computational demand (see Note 3), a minimal number of atoms is included in each simulation.

smog_extract, included as a part of the SMOG 2 package, can be used to prepare a structure-based model for a fraction of a larger assembly. Specifically, smog_extract will read a set of Gromacs-formatted force field files and generate new files that describe a subset of the original atoms. In doing so, it ensures that the truncated system provides an identical energetic representation of the preserved atoms. Using smog_extract is also simpler, in terms of user intervention, than removing atoms from the PDB file and then using SMOG 2 to generate a new model. For example, if one were to use the latter approach, the prepared PDB file would still need to conform to the defined templates (e.g. every atom defined in each residue would need to be present). In addition, the force field could be perturbed by the use of a truncated PDB (e.g. scaling of energetics is system dependent [13] and the generated contact map could be impacted by the absence of atoms).

smog_extract processes input structure (.gro) and topology (.top) files describing the molecular system, and then produces the corresponding files for a truncated system. The list of atoms to be included in the truncated system is supplied as a Gromacs-formatted index file (.ndx). This list of selected atoms can be prepared using your choice of molecular visualization programs, such as VMD (See Note 4). A sample invocation of smog_extract would be:

~~~
>$ smog_extract *−* ffullsystem.top *−*gfullsyst m.gro *−*n truncated.ndx
~~~

A general concern when simulating a subsystem is that the removal of atoms may lead to artificial changes in molecular flexibility at the boundary of the truncated system. To address this, one may optionally instruct smog_extract to introduce a harmonic position restraint on every atom that has an interaction (bond, bond angle, dihedral, contact) removed during the truncation step. For this, the user simply needs to supply the flag -restraint <val>, where <val> is the energetic weight of the restraints. The restraint strength is in units of energy/nm^2^, where energy is in reduced units (See Note 5). As a side note, since reduced units are employed, it is important that one properly interprets the reported simulated timescale. For a discussion on time scale estimates, see Note 6.

### 3.3 Avoiding artificial boundary effects in truncated systems

One of the strengths of structure-based models is that they are able to provide descriptions of the overall flexibility of biomolecules that are consistent with experimental observations and more highly-detailed models. For example, the root mean squared fluctuations (rmsf) of each residue in the ribosome have been shown to be similar between SMOG models, explicit-solvent simulations and crystallographic B-factors [23]. Similarly, the structural fluctuations of single molecules are also often consistent between SMOG and explicit-solvent models [5, 24]. Since the scale of these fluctuations can directly influence the kinetics of other conformational processes and free-energy barriers [10, 22], an accurate representation of flexibility is important when studying any molecular assembly.

By truncating a molecular system using smog_extract (see Section 3.2), the mobility of the atoms at the system boundary are likely to be perturbed. To avoid introducing these artificial effects, it is necessary that one tunes the strength of the atomic restraints imposed on the boundary atoms. Below, we describe a fluctuation-matching protocol that has been applied to study the ribosome [10]. In this approach, heterogeneous isotropic spatial restraints are refined, such that the dynamics in the truncated systems are consistent with the full assembly.

To apply fluctuation matching techniques, the user must establish reference values for the mobility of each atom and then introduce atomic restraints in the truncated system. To define the reference fluctuations, the following steps may be applied: First, generate a SMOG model and perform a simulation of the complete molecular system at a desired reference temperature. For tips on selecting a simulated temperature, see Note 7. Next, use smog_extract to generate a new set of top/gro files for the user-defined subset of atoms. It is necessary to use the -restraint option, in order to automatically introduce restraints on the boundary atoms. This will result in the position_restriants directive being added to the output .top file (see Listing 1).

Listing 1: Example .top file in which homogeneous isotropic position restraints are added by smog_extract.

~~~
[position _ restraints]
1 1 100.00 100.00 100.00
2 1 100.00 100.00 100.00
…
~~~

Finally, for the boundary atoms identified by smog_extract, calculate the rmsf values in the simulation of the full system. The Gromacs module rmsf may be used for this step.

~~~
>$ gmx rmsf *−* ftraj.xtc *−*s run.tpr *−*n boundary_ full.ndx *−*o rmsf.xvg
~~~

This will provide the rmsf value of each boundary atom, from which you can calculate the msf of each atom (msf_i_^ref^) and the average msf value of the boundary atoms <msf>^ref^. These will serve as the reference values during refinement.

After establishing a set of reference msf values for the boundary atoms, one needs to iteratively perform simulations of the truncated system and update the position_restraints section of the truncated .top file. The specific steps are:

1. Perform a simulation of the truncated system using the truncated .top file. Note: Since the potential energy is not translationally invariant when position restraints are included, it is important that the center of mass velocity is *not* removed during the simulation. That is, include comm_mode=none in the .mdp file.
2. Calculate the rmsf values of the boundary atoms.

~~~
>$ gmx rmsf *−* f traj.xtc *−*s run.tpr *−*n boundary_truncated.ndx *−* nofit
~~~ From this, calculate the mean msf value for the truncated system <msf>^trunc^.
3. Rescale the weight of every position restraint (listed in the top file) by the factor <msf>^trunc^/<msf>^ref^.
4. Return to step 1.

The above steps should be repeated until <msf>^trunc^/<msf>^ref^∼1, at which point the average msf values in the truncated system are consistent with the full system. However, this global rescaling does not ensure that the mobility of each atom is consistent. To further improve agreement between the truncated and full systems, the above steps may be continued, though in the subsequent iterations each atomic restraint should be rescaled individually by msf_i_^trunc^/msf_i_^ref^. After multiple iterations, one will obtain a topology file in which heterogeneous restraints are present (Listing 2).

Listing 2: Example .top file specifying isotropic position restraints.

~~~
[position _restraints]
1 1 154.338 154.338 154.338
2 1 122.95 122.95 122.95
…
~~~

It should be noted that more sophisticated refinement algorithms are available. For example, the method of Savelyev and Papoian follows a similar sequence of steps, though the updated values of the parameters (restraints) account for the possibility of coupling between restrained atoms [25]. In addition, the described approach may be extended to include anisotropic restraints [22]. For an example of the effectiveness of the approach, see Fig. 1.

**Figure 1:**
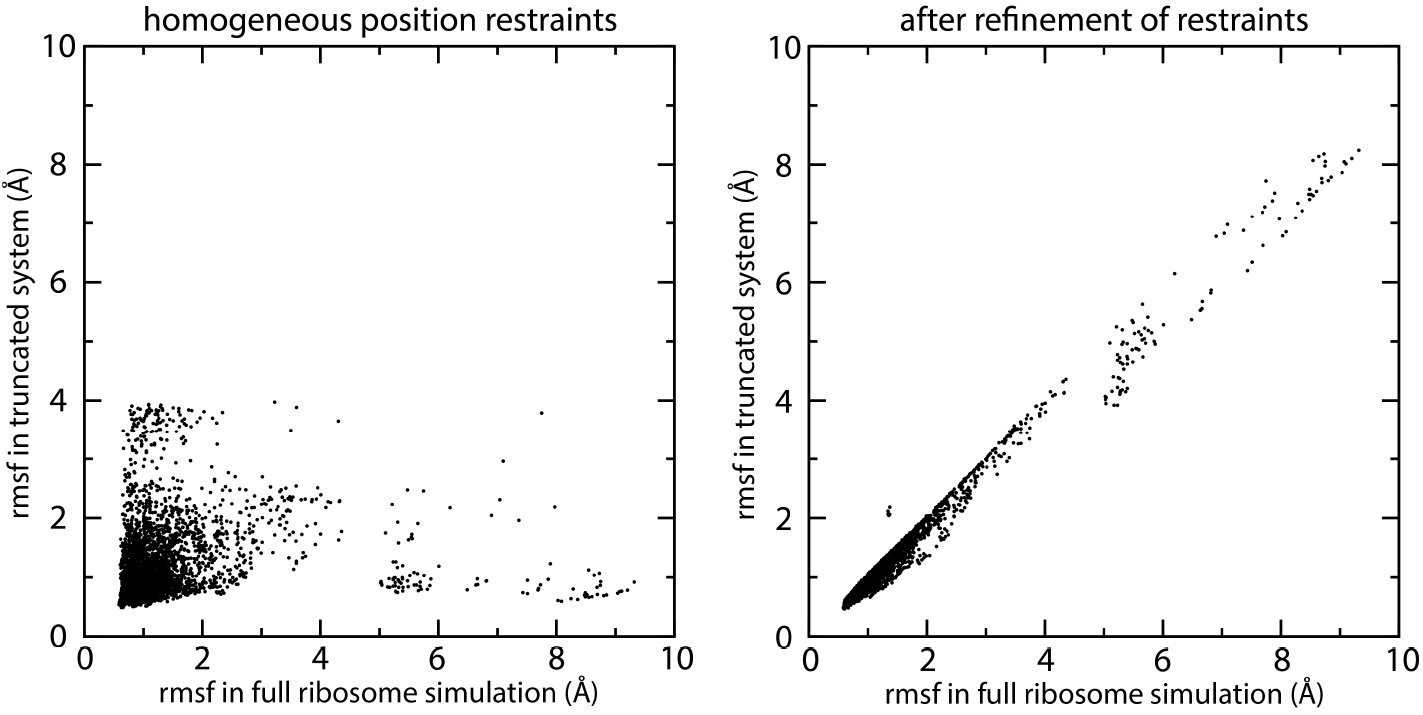
Comparison of rmsf values obtained for simulations of the ribosome and a truncated subset of ribosomal atoms. Left) Atomic rmsf values obtained in a simulation of a complete ribosome, compared to the values obtained in a truncated system in which homogeneous isotropic position restraints are applied. As expected, there is relatively poor agreement when a homogeneous value of the restraint is used for all atoms. Right) Comparison of rmsf values obtained after refinement of isotropic [10] and then anisotropic position restraints [22]. After refinement, the rmsf values have a correlation coefficient of 0.99 and a mean squared deviation of 0.03 Å^2^.

### 3.4 Extending structure-based models to include non-standard residues

The default SMOG 2 templates provide definitions of common amino acid and nucleic acid residues. However, there are many assemblies that are influenced by post-transcriptional and post-translational modifications, and large-scale conformational changes are often associated with ligand binding/release. To include these molecular features in SMOG models, one needs to define additional residue types. Here, we use phosphothreonine (TPO) as an example for how to add an amino acid definition to the SMOG 2 template files. For this example, we will demonstrate how to modify the standard all-atom structure-based model [5] that is distribute with SMOG 2. Since we are simply adding another amino acid (as opposed to adding inorganic molecules, or prosthetic groups), one only needs to introduce the following modifications to the biomolecular structure file (SBM_AA/AA-whitford09.bif):

1. Place a residue tag within the residues element (Listing 3). The residue tag requires at least two attributes: name (TPO) and type (amino). The residue name must be identical to the name appearing in the PDB file.

##### Listing 3: Amino acid residue section of .bif file

~~~
<!-- PHOSPHOTHREONINE -->
     < residue name=" TPO " residueType =" amino ">
        < atoms >
        </ atoms >
        < bonds >
        </ bonds >
        < impropers >
        </ impropers >
     </ residue >
<!-- THREONINE - TERMINAL -->
~~~
2. Define each atom in the residue by providing atom child elements within the atoms element. Similar to the residue name, the atom names must match the convention used in the input PDB file. Since we are extending the standard SMOG model, where all energetic parameters are homogeneous (e.g. all bonds have the same spring constant), the bType, nbType and pairType are given the same common values of B_1, NB_1 and P_1 (Listing 4). For instructions on how to include heterogeneous energetic parameters, consult the user’s manual.

##### Listing 4: Adding the atoms section to the residue structure

~~~
<! -- PHOSPHOTHREONINE -->
     <residue name =" TPO " residueType =" amino ">
     < atoms >
        < atom bType=" B_1 " nb Type=" NB_1 " pairType=" P_1 "> N</ atom >
        < atom bType=" B_1 " nb Type=" NB_1 " pairType=" P_1 "> CA </ atom >
        < atom bType=" B_1 " nb Type=" NB_1 " pairType=" P_1 "> CB </ atom >
        < atom bType=" B_1 " nb Type=" NB_1 " pairType=" P_1 "> CG2 </ atom >
        < atom bType=" B_1 " nb Type=" NB_1 " pairType=" P_1 "> OG1 </ atom >
        < atom bType=" B_1 " nb Type=" NB_1 " pairType=" P_1 "> P</ atom >
        < atom bType=" B_1 " nb Type=" NB_1 " pairType=" P_1 "> O1P </ atom >
        < atom bType=" B_1 " nb Type=" NB_1 " pairType=" P_1 "> O2P </ atom >
        < atom bType=" B_1 " nb Type=" NB_1 " pairType=" P_1 "> O3P </ atom >
        < atom bType=" B_1 " nb Type=" NB_1 " pairType=" P_1 "> C</ atom >
        < atom bType=" B_1 " nb Type=" NB_1 " pairType=" P_1 "> O</ atom >
     </ atoms >
     < bonds >
     </ bonds >
     < impropers >
     </ impropers >
    </ residue >
~~~
3. Insert definitions for the chemical bonds within the bonds element.

##### Listing 5: Adding the bonds to the residue definition

~~~
     < bonds >
<!-- BACKBONE -->
     < bond energy Group =" bb_a">
        < atom > N</ atom >
        < atom > CA </ atom >
     </ bond >
     < bond energy Group =" bb_a">
        < atom > CA </ atom >
        < atom > C</ atom >
     </ bond >
     < bond energy Group =" bb_a">
        < atom > C</ atom >
        < atom > O</ atom >
     </ bond >
<!-- FUNCTIONAL GROUP -->
     < bond energy Group =" sc_a">
        < atom > CA </ atom >
        < atom > CB </ atom >
     </ bond >
     < bond energy Group =" sc_a">
        < atom > CB </ atom >
        < atom > OG1 </ atom >
     </ bond >
     < bond energy Group =" sc_a">
        < atom > CB </ atom >
        < atom > CG2 </ atom >
     </ bond >
<!-- ADDITIONAL BONDS FOR THE PHOSPHATE GROUP -->
     < bond energy Group =" sc_a">
        < atom > OG1 </ atom >
        < atom > P</ atom >
                </ bond >
     < bond energy Group =" sc_a">
        < atom > P</ atom >
        < atom > O1P </ atom >
     </ bond >
     < bond energy Group =" sc_a">
        < atom > P</ atom >
        < atom > O2P </ atom >
     </ bond >
     < bond energy Group =" sc_a">
        < atom > P</ atom >
        < atom > O3P </ atom >
     </ bond >
     </ bonds >
~~~ Each bond element defines a single bond between two atoms. A required attribute of each bond is the energyGroup attribute, which indicates how to define dihedral interactions about the bond. In the default all-atom model, bb_a (sc_a) indicates that the bond is part of the protein backbone (side chain) and that the associated dihedral should be given cosine potentials.
4. Define any improper dihedral angles. An improper dihedral is used to ensure chirality about an atom for which not all bonded atoms are explicitly represented (e.g. due to the removal of hydrogen atoms), or to ensure that trigonal planar covalent geometry is maintained. Each improper dihedral associated with a residue should be listed within an improper element. For TPO, there are two such dihedrals: CB-CA-C-N and CA-CB-OG1-CG2.

##### Listing 6: Adding the improper dihedral section to the residue structure

~~~
    < impropers >
       < improper >
          < atom > CB </ atom >
          < atom > CA </ atom >
          < atom > C</ atom >
          < atom > N</ atom >
     </ improper >
     < improper >
          < atom > CA </ atom >
          < atom > CB </ atom >
          < atom > OG1 </ atom >
          < atom > CG2 </ atom >
     </ improper >
     </ impropers >
~~~

### 3.5 Including electrostatics in structure-based models

As discussed above, one of the key features of SMOG 2 is that it allows the user to adjust force field definitions without requiring source-code modifications. One way in which these models are often extended is to include an explicit representation of electrostatic interactions [8, 9, 10]. Here, we describe multiple ways in which these models may be extended to include electrostatic interactions and ionic effects.

#### 3.5.1 Assigning charges

There are two methods by which a user may define charges within the SMOG 2 framework. First, it is possible to define atom types that carry specific charges. The second method is to override atom type definitions and provide charge definitions for specific atoms within a defined residue.

##### Adding charges by atom type

It is common in classical mechanics force fields for one to provide identical energetic parameters for many chemically-similar atoms. For example, one may assign the same parameters (mass, charge, vdW) to every backbone P atom in RNA. To implement this in SMOG 2, one needs to modify the .nb and and .bif template files. The first step is to use the nonbond element to define a new atom type. In the example below (Listing 7) the nbType NB_P will be used to describe P atoms.

##### Listing 7: Modifying .nb file to change the charge and mass

~~~
<!-- GENERAL NONBONDS -->
< nonbond mass=" 1.00 " charge=" 0.000 " ptype=" A" c6 =" 0.0 " c12 =" 5.96046 e-9 ">
     < nb Type > NB_1 </ nb Type >
</ nonbond >
< nonbond mass=" 2.50 " charge=" -1.000 " ptype=" A" c6 =" 0.0 " c12 =" 5.96046 e-9 ">
     < nb Type > NB_P </ nb Type >
</ nonbond >
~~~

In this example, the NB_P type is defined to be of mass 2.5, charge of -1 and have only a repulsive non-specific vdW parameter. The second step is to use the nbType type within an atom element of a residue (e.g. Listing 4).

After modifying the template files to define a new atom type, these templates may be used with SMOG 2. After running SMOG 2, if an atom of the newly-defined type is present in the molecular system, then its definition will appear under the atomtypes directive (Listing 8) in the generate .top file.

Listing 8: Charge information shown in .top file

~~~
[atomtypes]
;   name   mass   charge   ptype  c6     c12
NB_1   1.0000   0.000000   A 0 . 00000 e+00     5.96046 e*−*09
NB_P   2.5000   *−*1.000000   A  0.00000 e+00     5.96046 e*−*09
~~~

##### Adding charges to residue definitions

While it is possible to always use nbType definitions to assign charges, it is sometimes more convenient to define charges for specific atoms within a particular residue. For example, one may add a charge to the P atom of each Adenine residue by adding the charge attribute to the atom element P for residue A. It should be noted that explicit assignment of charge to specific atoms will supersede any charge assignments based on the nbType. For example, if the P atom were given nbType of NB_1, which is defined to have charge 0, the explicit attribute charge="-1" would override this value. This type of assignment will result in charges appearing on specific atoms under the atoms directive of the .top file.

#### 3.5.2 Modeling monovalent and divalent ions

The dynamics of many biomolecular assemblies, especially those containing RNA (e.g. the ribosome and spliceosome), are strongly influenced by the presence of ions. Ions may bind to assemblies and contribute to structural stability, or the local environment of diffuse ions may lead to non-linear electrostatic screening effects. Depending on what aspects of ion dynamics you would like to study, there are multiple strategies for defining ions in structure-based models. Here, we describe multiple SMOG tools that can facilitate the study of ionic effects in biomolecular assemblies. Before discussing the technical aspects of introducing ions, it is important to note that the user is ultimately responsible for calibrating the most appropriate scale and functional form of the electrostatic interactions. While there are some general guidelines for calibrating energy scales, such considerations must be applied on a per-model basis (see Note 8).

##### Implicit treatment of ions

To simulate a system in which monovalent ions are treated implicitly, one may introduce electrostitics via a screened Debye-Hückel (DH) potential. To accomplish this, one needs to generate a look-up table that defines the functional form of the desired electrostatic potential. This table is then provided as input to Gromacs. For this step, SMOG 2 provides the tool smog_tablegen, which may be invoked with the following flags:

~~~
>$ smog_tablegen *−*M <M> *−*N <N> *−* i c <ion conc .> *−*sd <elec . switch dist.> \\
*−*sc <elec.truncate dist.> *−* tl <table length > *−*table <output name>
~~~

Here, <M> and <N> denote the exponents of the attractive and repulsive non-bonded interactions, respectively. If M and N are not provided, default values of 6 and 12 are used. <ion conc.> is the desired effective monovalent ion concentration, which determines the Debye screening length, as implemented previous [8]. Finally, in order to ensure continuous first derivatives, a fourth-order polynomial is added to the force over the distance range <elec. switch dist.> to <elec. truncate dist.> (nm).

Employing the DH potential in a specific simulation requires minor changes to both the grompp and mdrun steps. Before running grompp, the mdp file must define coulombtype=User and vdwtype=User. When running the simulation with mdrun, the user has to indicate where the table file is located:

~~~
>$ gmx mdrun *−*s run . tpr *−*noddcheck *−*table table. xvg *−*table p table. xvg
~~~

In this example, table.xvg is located in the current working directory.

##### Explicit treatment of ions

In addition to implicitly accounting for monovalent ions, SMOG 2 also supports explicit ion models. Explicit ions may be treated as structural (bound), or diffuse/bulk. For these two representations, different steps should be followed.

###### Bound/Structural ions

One way in which to describe bound ions is to treat them as part of the biomolecular structure. For example, crystallographic models of the ribosome often include “structural” Mg^2+^ ions. Since the residence time of these ions is much longer than accessible simulation times, it is appropriate to describe these ions as being permanently bound to the biomolecular complex. For this representation, SMOG 2 can read a PDB file in which ions are present and then include harmonic interactions between the ions and molecular system. Consistent with the general approach to defining structure-based models, each harmonic potential will have a minimum corresponding to the distance found in the provided PDB structure. In the default all-atom model, this type of interaction is defined for atom and residue type BMG ("Bound MG"). This treatment of ions is declared in the .nb and .bif template files. First, the BMG residue is defined in the .bif file (Listing 9).

##### Listing 9: Defining ions in the .bif file

~~~
            <!-- ION RESIDUES -->
<!-- Bound MG ions -->
     < residue name=" BMG " residueType =" ion" connect=" no" atom Count=" 0 ">
       < atoms >
          < atom bType=" B_1 " nb Type=" NB_1 " pairType=" BMG " bonds=" 0 "> BMG </ atom >
      </ atoms >
     </residue>
~~~

In this definition, one should notice that there are several additional attributes provided to the residue and atom elements. Specifically, the connect=“no” attribute instructs SMOG 2 to not attempt to include bonds between ions that are listed sequentially in the input PDB file. The next attribute to notice is atomCount=“0”, which instructs SMOG 2 to *not* account for ions when setting any energetic normalization conditions. Finally, bonds=“0” indicates that the ion has no covalent bonds. If this attribute is absent and a BMG atom appears in your PDB file, SMOG 2 will exit with an error, since it expects every atom to have at least one covalent bond. In addition to defining the BMG residue in the .bif file, the template files indicate that harmonic interactions should be included with BMG atoms (rather than the typical 6-12 potential for contacts). This is declared in the .nb file (Listing 10).

##### Listing 10: Defining the contact potential for bound BMG ions

~~~
   <contact func =" bond_type6 (?, 200) " contactGroup ="c">
      <pairType >BMG </ pairType >
      <pairType >*</ pairType >
   </ contact >
~~~

Here, the function type is bond_type6, which is defined as a harmonic, non-chemical, potential (see Gromacs manual for details). It should be noted that the user is at liberty to use any pair-wise potential to describe ion-biomolecule interactions. In the above discussion, we have used a harmonic representation as an example.

###### Diffuse/Bulk ions

In addition to structural ions, the precise local concentration of freely diffusing ions can strongly influence the kinetics of large-scale conformational processes. If one is aiming to study the role of diffuse ions with structure-based models, then it is necessary to add additional ions that are not present in the input PDB structure. Within the SMOG framework, after generating a force field for the biomolecule using the executable smog2, additional ion definitions must be present in the .top and .gro files. The tool smog_ions was written for the specific purpose of adding a user-defined number of ions to a structure-based model force field. In addition to providing the ion name and number of desired ions, the user must also specify the charge, mass, and c12 parameter that defines its excluded volume (an attractive c6 term is optional). Note that each call of smog_ions can only add a single ion type. Thus, if you would like to add multiple ion species (e.g. K^+^ and Cl^*−*^), it would be necessary to make repeated calls to smog_ions. After running smog_ions, the user will have a .top file describing the composite biomolecule-ion system, as well as a new coordinate file that will have randomly placed ions (e.g. Fig. 2)

**Figure 2:**
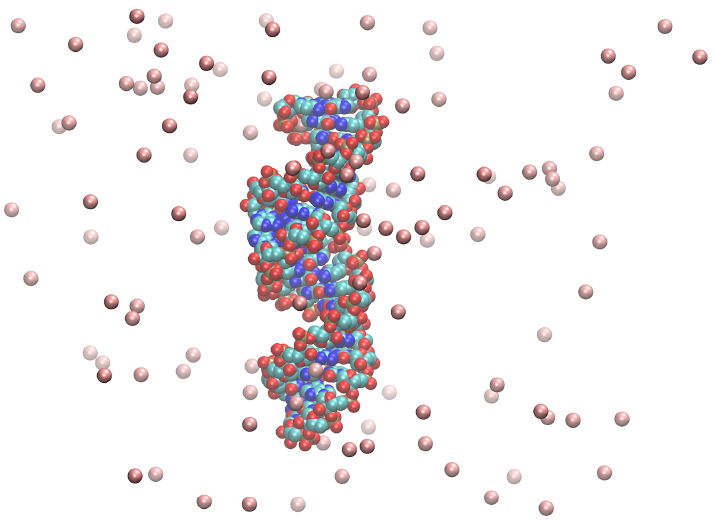
K^+^ ions (pink beads) added to an RNA model using smog_ions

## 4 Notes

1. Some common processing errors:

- Missing atoms **–** Structures with insufficient resolution may be missing atoms because the local electron density didn’t allow for its determination. If these regions are in disordered loops, a simple solution is simply to edit these residues’ names to ALA and remove any side chain atoms beyond the CB. The reasoning would be that any native contacts that may be excluded would likely have been an artifact of the crystallization conditions. If the missing atoms are not in loops, an appropriate strategy may be to use homology modeling software to insert the missing atoms, e.g. SWISS-MODEL [26].
- Non-canonical residues, such as MSE **–** X-ray crystallography is often aided by small perturbations such as selenomethionine (MSE) instead of methionine (MET). In the case of MSE, the solution is either a. to add MSE to the .bif by duplicating MET and changing S atom to SE, or b. to edit the PDB by renaming MSE to MET. If the bonded structure is not exactly identical to one of the residues in the .bif file, another strategy would be to generate a homology model to restore the native sequence. This model would then be used as input to SMOG 2.
- Missing residues **–** Often loop regions will be missing due to disorder in crystal structures. One solution is to insert TER lines between breaks in the protein sequence. However, one issue with this approach is that it requires that the simulated temperature to be sufficiently low that the now disconnected chains do not dissociate. Homology modeling software can be used to insert the missing residues, but this raises the question of whether to add native contacts for the disordered region. To automatically ignore contacts for these disordered regions via the SMOG 2 templates: a. duplicate all the residues in the .bif with names (e.g. ALA->ALAD), b. change all the pair types for atoms in these new residues to a new type (e.g. P_1 -> P_D), c. add a rule in the .nb for contacts between anything (type *) and P_D with func="contact_free()".
2. When running protein folding simulations, take care to run the simulation with a sufficiently large box. The xyz dimensions of the box are denoted in the last line of the .gro file. The protein should never interact with itself through the periodic boundaries. Take note that Gromacs tabulated pair potentials (the Lennard-Jones native contacts) are neglected for the remainder of a simulation if the distance between a naive pair exceeds the length of the table. This length is initialized as the largest cutoff distance, rvdw or rcoulomb. Since unfolded protein native pairs are far apart, setting table-extension in the .mdp to half the box diagonal can ensure that no pairs are inadvertently neglected.
3. Even though structure-based models are less computationally demanding than explicit-solvent models, simulations of large assemblies can require substantial resources. Fortunately, most modern MD engines exhibit strong scaling, such that many cores may be used for a single simulation. For a eukaryotic ribosome (250,000 atoms), Gromacs v4.6.3 was shown to scale to more than 1,000 compute cores [13]. In a previous study where only 1/6 of the ribosome was explicitly represented [27], a single trajectory required 128 cores for over 4 months. As a guide for expected performance, we have obtained more than 50,000,000 timesteps per day using 28 compute cores for a system of *≈*28,000 atoms [22].
4. When using smog_extract, the TkConsole of VMD can be a very helpful tool for generating the list of atoms to be included in the truncated system. In the example below, one can select a rectangular box of atoms and write the indices to the file truncatedAtoms.ndx:

~~~
**set** p0 [atomselect 0“ (x>85)and (x<140) and (y>105) and (y<155) and (z >140) and (z <180)”]
**set file** [**open** “truncated Atoms.ndx” w]
**puts** $ file [$p0 **get** serial]
**close** $ file
~~~ To fully adhere to .ndx file format, the user simply needs to add an atom group declaration (e.g. [group1]) to the first line of the .ndx file. Note that the above example uses the keyword “serial” (starts at 1), rather than “index” (start at 0).
5. SMOG 2 topology files are written in reduced units. The length unit is the same as Gromacs: nanometers. The mass of each bead is 1. In the all-atom model this corresponds to a mass unit of ~ 12 amu. In the C-alpha model, each residue is a single bead, which, given *≈* 8 heavy atoms per bead, corresponds to a mass unit of ~ 100 amu. In principle, the correct heterogeneous masses could be written to the topology, but this would only have a small effect on the kinetics, and thermodynamics is unaffected. A Gromacs .mdp expects the temperature to be provided in Kelvin, where a value of Boltzmann’s constant ._B_ of 0.00831451 kJ/mol/K is used internally. Thus, in a Gromacs .mdp file, a temperature of 120.3 K corresponds to a reduced temperature of 1. When adding additional energetic terms that have empirical units with the effective energetics in structure based models, it is often useful to express both energy scales in terms of k_B_T.
6. Estimating time scales in simplified and/or coarse-grained simulations can be a tricky business. In the structure-based model, speed-up relative to all-atom explicit-solvent models comes from three effects: a. unfrustrated energetics, b. low viscosity, c. coarse graining. In the field of protein folding, the first two effects are known as internal and external friction, respectively [28]. The external friction is formally absent because of the lack of solvent, but is re-introduced to a small degree through the use of Langevin dynamics protocols. The lack of non-native interactions in the structure-based model smooths the energy landscape and reduces the internal friction. Coarse-graining smooths the internal friction as well, by both simplifying and softening the interactions. Of course, coarse-graining also provides an obvious algorithmic speed-up by requiring less computation, but this is associated with the time scale. Taken together, for protein folding, the time scale is estimated to be increased by a factor of 1000-10000 [29]. Simple diffusion limited processes like molecular encounter in aggregation will be most dependent on the residual viscosity controlled with the thermostat and, thus, care must be taken when studying the kinetics of systems that involve multiple physical processes, e.g. coupled binding and folding of a homodimer.
7. Since reduced units are used in SMOG models, it is important to choose an appropriate simulated temperature. To determine the the proper value of the reduced temperature, one typically compares atomic rmsf values between a SMOG model and explicit-solvent simulations, or experimental B-factors. When using the all-atom SMOG model, a reduced Gromacs temperature of approximately 60-80 typically corresponds to a temperature of around 300 K in explicit-solvent simulations [24]. When studying biomolecular folding, an alternate strategy for calibrating temperature is to describe the system in terms of the folding temperature ._f_ . That is, by equating an experimental observable ._f_ with a simulation ._f_ can give a helpful point of reference. As a final note, it is important to recognize that linear extrapolation of temperature for any molecular mechanics model is not likely to be reliable, since the solvent introduces strong non-linear effects, e.g. cold denaturation and boiling.
8. After calibrating the temperature in the model, additional energetic terms can be included in the model and calibrated by thermal energy matching. Two examples that have been explored with SMOG models are pulling forces [30, 31] and electrostatic forces [10]. A correspondence between a experimental temperature *T*_exp_ = 300K and simulation temperature *T*_sim_ = 0.9 allowed for thermal energy matching, where *k*_B_*T*_exp_ = 2.5 kJ/mol = 4.2 pN . nm = *k*_B_*T*_sim_ = 0.9ϵ = 0.9[F]. nm. Thus, a reasonable initial estimate of the reduced force unit [F] is 4.2/0.9 pN = 4.7 pN = [F]. As a tip, for these types of comparisons, the strength of the Coulomb force may be adjusted by scaling the effective dielectric constant, which is set by epsilon_r in the mdp file. Given the same temperature calibration as above, one could rescale the dielectric constant by a factor of 4.2.0.9 = 4.7, where the dielectric for water 80 would be scaled to 80 *×* 4.7 = 376 → epsilon_r.

## 5 Acknowledgements

This work was supported in part by an NSF CAREER Award (Grant MCB-1350312). JKN is a Humboldt Postdoctoral Fellow. UM acknowledges support as a John Simon Guggneheim Memorial Foundation fellowship. We acknowledge generous support provided Northeastern University Discovery Cluster.

